# SCN9A gene therapy: Novel mechanism to inhibit cellular senescence in astrocytes *in vitro*

**DOI:** 10.1101/2023.07.24.549756

**Authors:** Divyash Shah, Donna Leonardi, Alyssa Waldron

**Affiliations:** University of Pennsylvania 200 Hackensack Avenue, Hackensack, NJ 07601 USA; Bergen County Academies 200 Hackensack Avenue, Hackensack, NJ 07601 USA

**Keywords:** aging, senescence, neurodegeneration, astrocytes, Alzheimer’s

## Abstract

Cellular aging, also known as senescence, is a form of proliferation arrest that occurs in cells with age. In individuals, this process occurs due to telomere degradation and consequent dysfunction. In the nervous system, senescence of astrocytes, the most common neuronal support cells, has been associated with inflammation and forms of neurodegeneration associated with various disease pathologies. Thus, studying senescence may be a unique approach to understanding brain and aging and consequential diseases as well. Studies have shown that transcription downregulation of SCN9A led to reversal of senescence in epithelial cells. So, this study explores the role of SCN9A in astrocytes.

*In silico* patient gene analysis reveals multiple significant pathways associated with neuronal aging including ion channel regulation. Subsequent analysis shows the downregulation of SCN9A is associated with genes in the neuronal senescence pathway. *In vitro* studies showed that astrocytes with knockdowned SCN9A did not undergo senescence as significantly as regular astrocytes. Furthermore, protein analysis presented a novel role for SCN9A in the astrocyte senescence pathway and an association with preventing DNA leakage. This study suggests SCN9A plays a large role in neurodegeneration and it should be studied in reversing senescence and even treatment plans for brain aging.

## 1. Introduction

The current model for understanding cellular aging is a form of proliferation arrest known as senescence.^[1]^ Senescence can be activated in cells via numerous external factors including DNA-damage, telomere shortening, or oxidative stress.^[1]^ The most popularly accepted senescence pathway is based on the involvement of cell cycle regulators including p21/p53.^[2]^ External stresses, most notably DNA damage, can induce activation of a cascade starting with inducing Cyclin-dependent kinase inhibitors (CDKIs) which eventually leads to the expression of p16/p21.^[1]^ These proteins in conjunction with various CDKIs ultimately prevent the phosphorylation of Retinoblastoma tumor suppressor (Rb) which develops into cell cycle arrest or senescence via the Rb/E2F pathway.^[1,2]^ As cells become senescent, they not only lose their normal functionality but also develop two key factors: the senescence-associated secretory phenotype (SASP) and Senescence-associated beta-galactosidase activity (SA-β-gal).^[2]^ These factors have been studied extensively with their correlation with age-related changes and diseases. Specifically, their presence in the nervous system has suggested a correlation with neuroinflammation and decline in brain function associated with IL-1 and IL-6. Neuroinflammation has also been studied with the newly developed DAMP hypothesis. In an *in vivo* mouse model, this decline is most notably present in synaptic function, thus indicating a connection between senescence cells and the neurodegeneration that occurs in the brain with age.^[3]^

The most common cells of interest in relation to brain aging are astrocytes, a type of glial cell that makes up a significant portion of the cell population in the central nervous system (CNS). Astrocytes have a critical role in neuronal energy metabolism via the synthesis of glutamate and γ-aminobutyric acid (GABA).^[4]^ Furthermore, astrocytes have enzymes that uniquely target neurons. As well, rain aging and various pathologies of age-related diseases have been linked with alterations in astrocyte function.^[4]^ These characteristics alongside astrocyte ability to modify neurotransmitter expression are the reasons astrocytes are of interest when discussing brain changes. Furthermore, while neurons naturally enter proliferation arrest after maturity, glial cells and astrocytes do not. This poses an interesting case to study especially because astrocytes are the support system for primary neurons and assist them in action potential signaling and management of synapses.^[5]^ *In vitro* models have shown that astrocyte function is dependent on their electrophysical properties which contributed to their hyperpolarization.^[6]^ This electric potential is the result of the various membrane-bound voltage-gated ion channels present on the surface of astrocytes.^[6]^ These ion channels contribute to astrocyte function and ion channel dysfunction has even been linked to neuroinflammation through a process known as reactive astrogliosis.^[5,7]^ Therefore, damage to glial cells would lead to decline in brain metabolism and function as a whole. When looking at the relationship between ion channels and aging, various genes that encode for ion channels have been associated with senescence in the past including SCN9A which encodes the sodium-voltage gated ion channel Nav 1.7.^[8]^ “Stressed” astrocytes may also exhibit damage-associated molecular pattern molecules (DAMPs) during senescence.^[9]^ As the DAMP hypothesis suggests biomarkers associated with aging and age-related disorders, studying their appearance may be a way to measure astrocyte senescence.^[10]^

Thus, a novel question presents itself in using our understanding of the astrocyte senescence pathway to reverse senescence *in vitro*. While initial understanding of senescence assumed that it was an irreversible process, current research has presented a new hypothesis that suggests that senescent cells can reenter the cell cycle under appropriate stimulus. However, the question of how to reverse senescence whether *in vitro* or *in vivo* still remains the larger unanswered issue. One gene identified in HEC-™ cells was SCN9A; Knockout of SCN9A was shown to prevent cellular senescence in epithelial cells which suggests a promising role for the gene in other cell types with similar phenotypes. With implications of various diseases including Alzheimer’s and Parkinson’s being associated with senescence, the importance of presenting a new understanding of astrocyte senescence is crucial.

Here, using both *in silico* and *in vitro* methodology, we assess if the role of SCN9A inhibition in astrocytes would prevent the cell from entering the state of senescence. This study hopes to confirm SCN9A’s role in cellular senescence and suggest the implications of SCN9A in the astrocyte model (4). Further, patient gene expression profiles were examined to highlight significant genes that play a role in neuroinflammation when correlated with aging. Further, *in vitro* model studies the role of SCN9A knockout in relation to key senescence markers and presents a potential correlation between senescence and DNA leakage.

## 2. Results

### 2.1 Identification of DEGs associated with aging in cortical neurons

In order to understand the molecular and cellular changes associated with aging in the Central Nervous System (CNS), gene expression dataset, GSE53890, was downloaded from GEO database.^[11]^ The GEO2R method was used to compare the expression profiles for cortical neurons in 41 patients grouped into two cohorts: age > 65 (n=21) and age < 65

(n=20). Differentially expressed genes (DEGs) were identified using commonly accepted significance (P < .05) and expression cutoffs of |logFC| ≥ 1. Results of GEO2R analysis identified 181 DEGs that were associated with aging in cortical neurons (**Figure 1**). 76 of these genes were upregulated in the older group (age > 65) and 105 genes were upregulated in the younger group (age < 65). These 181 DEGs were then analyzed in STRING to look for interactions between the various genes and their corresponding proteins.

**Figure 1.** Volcano plot of all differentially expressed genes (DEGs). Red dots represent upregulated genes in the older patient cohort with adj p<.05; blue dots represent downregulated genes with adj p<.05. The two black lines represent the logFC threshold and only genes with a |logFC| ≥ 1 were used for further analysis. Image produced in GEO2R.^[12]^

Following identification of DEGs, the online software STRING was used to construct a Protein-Protein Interaction (PPI) network using interactions between all 181 DEGs.^[13]^ The analysis resulted in a gene interaction had 138 active nodes with 263 edges and a PPI enrichment p=1.0e-16 (**Figure 2**).

**Figure 2.**
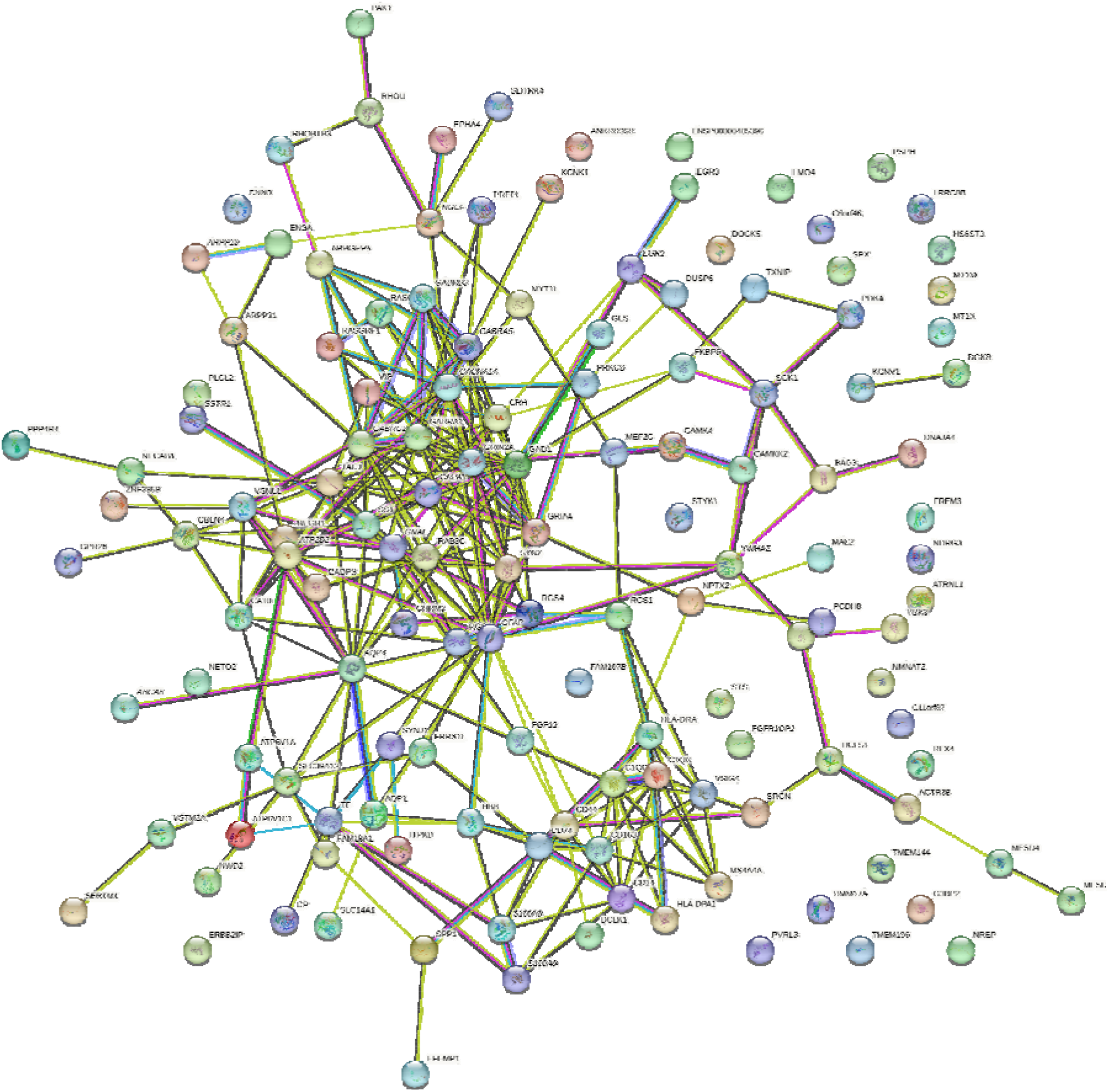
STRING Protein-Protein Interaction (PPI) map for DEGs in GSE53890. Each node represents a DEG and each line represents a potential interaction. PPI- *p=1.0e-16*. Image produced in STRING.^[13]^

Further Gene Ontology (GO) analysis, which looks at previously established pathways and compare to see how our DEGs are involved in these pathways, revealed that 34 of the 181 genes played the most significant role (strength >.9) in various molecular functions including neurotransmitter receptor activity, gated ion channels activity, and Neuroinflammation and glutamatergic signaling (**Table 1**).^[13]^ This suggests that the same genes responsible for neurotransmission and ligand-gated ion channel activity may play a significant role in or be significantly associated with neuronal aging processes.

**Table 1.** Results of STRING Gene Ontology analysis for DEGs in GSE53890 suggested various key molecular functions that were associated with the genes. These functions are thus presented to be associated with aging in the brain as well.

| Molecular Function | Count in Network | Strength | False Discovery Rate (FDR) |
| --- | --- | --- | --- |
| GABA-A receptor activity | 4 of 19 | 1.47 | 0.0063 |
| Transmitter-gated ion channel activity involved in regulation of postsynaptic membrane potential | 6 of 53 | 1.21 | 0.0057 |
| Postsynaptic neurotransmitter receptor activity | 7 of 70 | 1.15 | 0.0037 |
| Neuroinflammation and glutamatergic signaling | 9 of 140 | 0.96 | 0.00072 |
| Ligand-gated ion channel activity | 8 of 133 | 0.93 | 0.0057 |

### 2.2 Identification of DEGs associated with SCN9A Knockdown

Since GO analysis from the previous analysis suggested ion channel activity had a significant role in nervous system aging, analysis of SCN9A, a gene encoding for the NaV1.7 ion channel, in senescence may be noteworthy to see if its role translates to astrocyte senescence. GEO dataset GSE110884 contains gene expression profiles of epithelial cells treated with shRNA SCN9A, an *in vitro* technique to knockout effect of SCN9A in order to prevent senescence.^[8]^ GEO2R analysis revealed 1675 DEGs when comparing the genomic profiles of cells with shSCN9A (treated) vs cells without (control) after 72-hrs of 4-hydroxytamoxifen (4-OHT) to induce senescence (**Figure 3**).

**Figure 3.** Volcano plot of all differentially expressed genes (DEGs) from GSE110884. Red dots represent upregulated genes in cells with SCN9A knocked out when treated with 4-OHT for 72hrs with adj p < .05; blue dots represent downregulated genes in same cell conditions with p < .05. The two black lines represent the logFC threshold and only genes with a |logFC| ≥ 1 were used for further analysis. Image produced in GEO2R.^[12]^

Differentially expressed genes, both upregulated and downregulated were then subject to subsequent analysis using Enrichir.^[14]^ Enrichment analysis suggested key biological processes associated with the knockdown of SCN9A to prevent senescence in cells. The most notable processes associated with the 1675 DEGs were DNA metabolic processes (GO:0006259) [p=2.4e-13], DNA replication (GO:0006260)[p=2.53e-13], centromere complex assembly (GO:0034508)[p=5.33e-13], DNA synthesis involved in DNA repair (GO: 0000731)[p=173e-10]. Analysis results suggest that key cell cycle pathways that are responsible for keeping the cell proliferating excrete the same proteins that were also potentially synthesized by treatment of shSCN9A.

### 2.3 Determining IC50 of D-galactose

In order to model the cellular reaction to senescence, an *in vitro* methodology must be created to induce cell-cycle arrest pressure. D-galactose is suggested to be extremely effective in triggering brain aging via some oxidative stress pathway. Previous studies have shown that using D-galactose could suppress cell viability and induce cellular senescence.^[15]^ Therefore, cultured Astrocyte Type III Clones (C8D30s) were treated with D-galactose to identify the optimal concentration to induce senescence. The IC-50 (half maximal inhibitory concentration) is considered the most appropriate concentration for D-Galactose since this is the ideal value to induce senescence under enough stress without consequently causing significant apoptosis.^[15]^ Thus, astrocytes were cultured with varying concentration of D-galactose (0-555mM) and cell viability was assessed after 48 hours (**Figure 4**).

**Figure 4.**
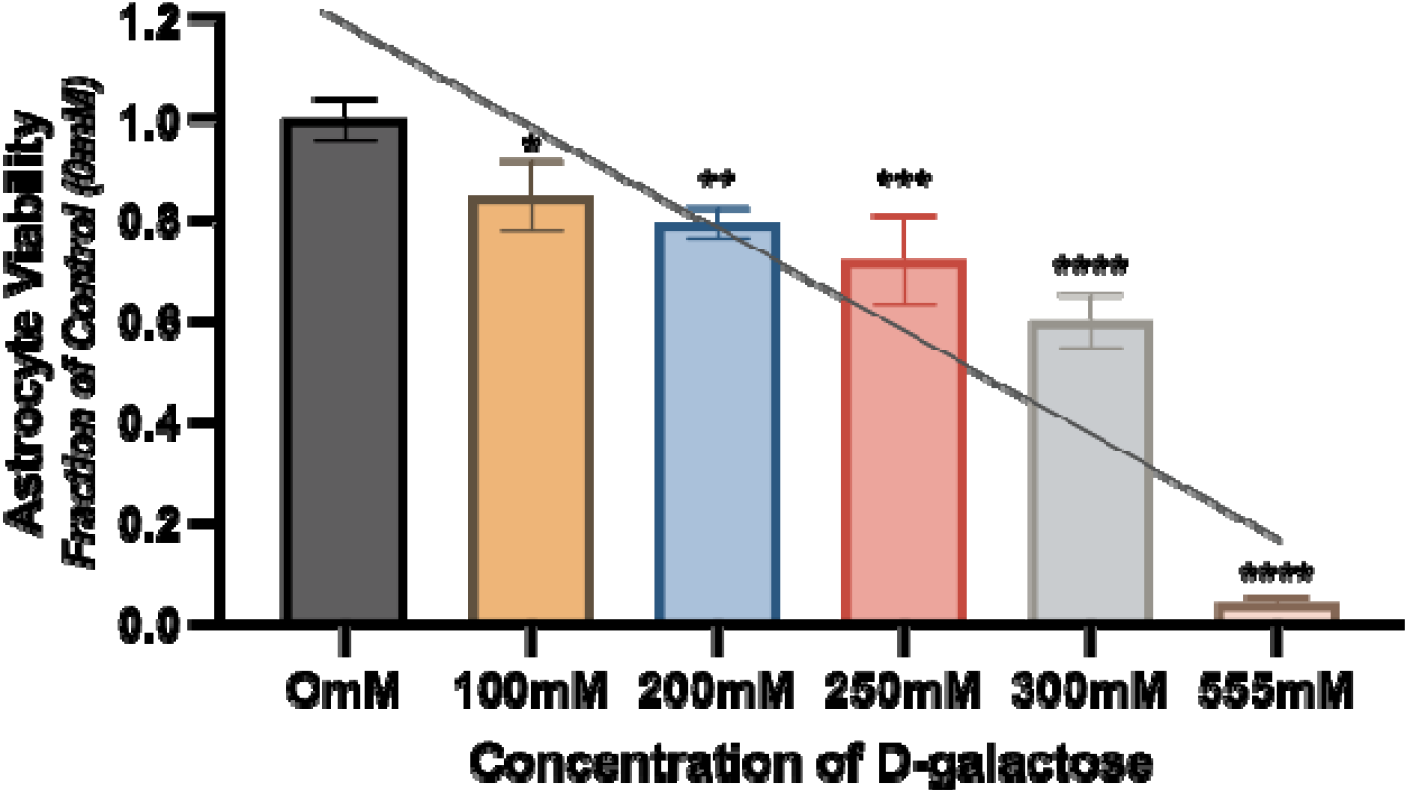
Varying concentration of D-galactose induces different responses in Astrocyte Type III Clones. Most notably, cell viability dropped off after 300mM. The IC-50 was determined, using a linear regression line, to be 250mM (R^2^=.806). *p<.05 and **p<.01 ***p<.001 ****p<.0001 for one way ANOVA with Dunnett’s multiple comparison test.

Most notably, treatments greater than 300mM significantly suppressed cell viability by more than 40% and even induced apoptosis in cells as noted by the presence of floating cells when viewed under light microscopy. Using the trendline seen in Figure 4, the IC50 was determined to be 250mM which would be used to induce senescence in cells in future experiments.

### 2.4 SCN9A knockout inhibits entrance into senescence in astrocytes after D-galactose exposure

To attempt to understand the role of knocking down SCN9A in preventing cells from entering senescence, astrocytes were transfected with SCN9A-targeting shRNA or scrambled RNA or control. Success of transfection was verified using fluorescence microscopy. Cells were treated with 250mM D-Galactose for 72 hours to induce senescence.

The most accepted marker of senescence is Senescence-associated-□-Galactosidase (SA-□*-*gal). Cells were stained and presence of SA-□*-*gal was analyzed using microscopy (**Figure 5**).

**Figure 5.**
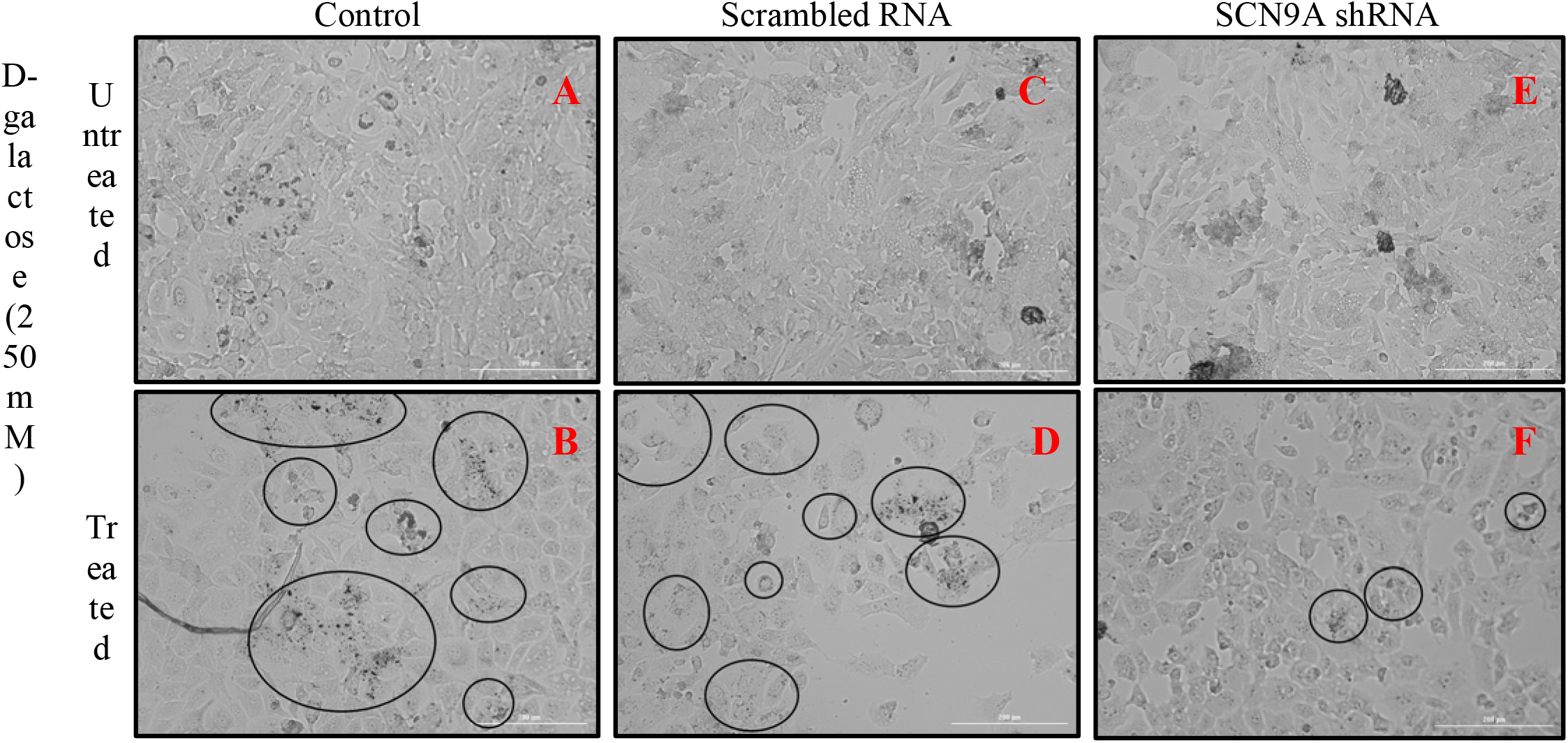
Loss of SCN9A in C8D30 prevented cells from entering D-galactose induced senescence. Cells of three conditions (Control, scrambled RNA, or SCN9A shRNA) were either treated with D-gal for 72 hours or left in the media. (A) Control cells appear to be proliferating normally with no change in morphology. (B) Cells after D-gal treatment appear to enter proliferation arrest as noted by lack of confluency and appearance of □*-Galactosidase* as indicated by annotated circles. (C) Scrambled RNA plasmid cells follow the same pattern as control cells as expected. (D) Scrambled RNA plasmid cells treated with D-gal also became senescent as noted by patchy cell clusters and appearance of □*-Galactosidase* in annotated cells. (E) Again, shSCN9A cells that were left untreated had no notable change as compared to control in either morphology or proliferation. (F) SCN9A knockdown cells also did not enter senescence as hypothesized. With the knockdown of SCN9A, all cells were not able to resist D-galactose stress as □*-Galactosidase* appeared in the staining of some cells (*annotated circles)* but majority of cells in this representative image did not enter proliferation arrest due to the SCN9A knockdown.

As expected, exposing astrocytes to D-galactose induced senescence led to changes in the morphology and clustering of cells(Figure 5B, 5D). Comparison of the staining of □*-Galactosidase*, a hallmark of senescence in astrocytes, of the untreated and treated astrocytes also supported induction of senescence. However, the majority of astrocytes containing the shSCN9A plasmid did not undergo senescence when they were exposed to D-galactose (Figure 5F). Noting the change in morphology that appears in scrambled and control cells under D-galactose, it is seen that the astrocytes treated with SCN9A shRNA were able to proliferate more effectively as the representative image suggests a more confluent image with shSCN9A (Figure 5B,5D,5F). The results here suggest that SCN9A has the potential to prevent astrocytes from entering senescence: a novel discovery.

To further verify microscopy results, cells were imaged under Texas Red And □*- Galactosidase* activity was measured at an optimal absorbance value (587 nm). In comparing the control cells to scrambled plasmid cells treated with D-galactose, there is a clear difference in fluorescence between the two images indicating a larger concentration of □*-Galactosidase* in treated cells corresponding to cells entering senescence(**Figure 6A**, 6C). This light intensity is quantified when images are converted into a gray-scale image. It is seen that scrambled RNA cells treated with D-galactose had a higher peak gray value close to 39,000 bits as compared to the control cells with a peak gray value of 33,000 bits which is supported by the images (Figure 6B, 6D).

**Figure 6.**
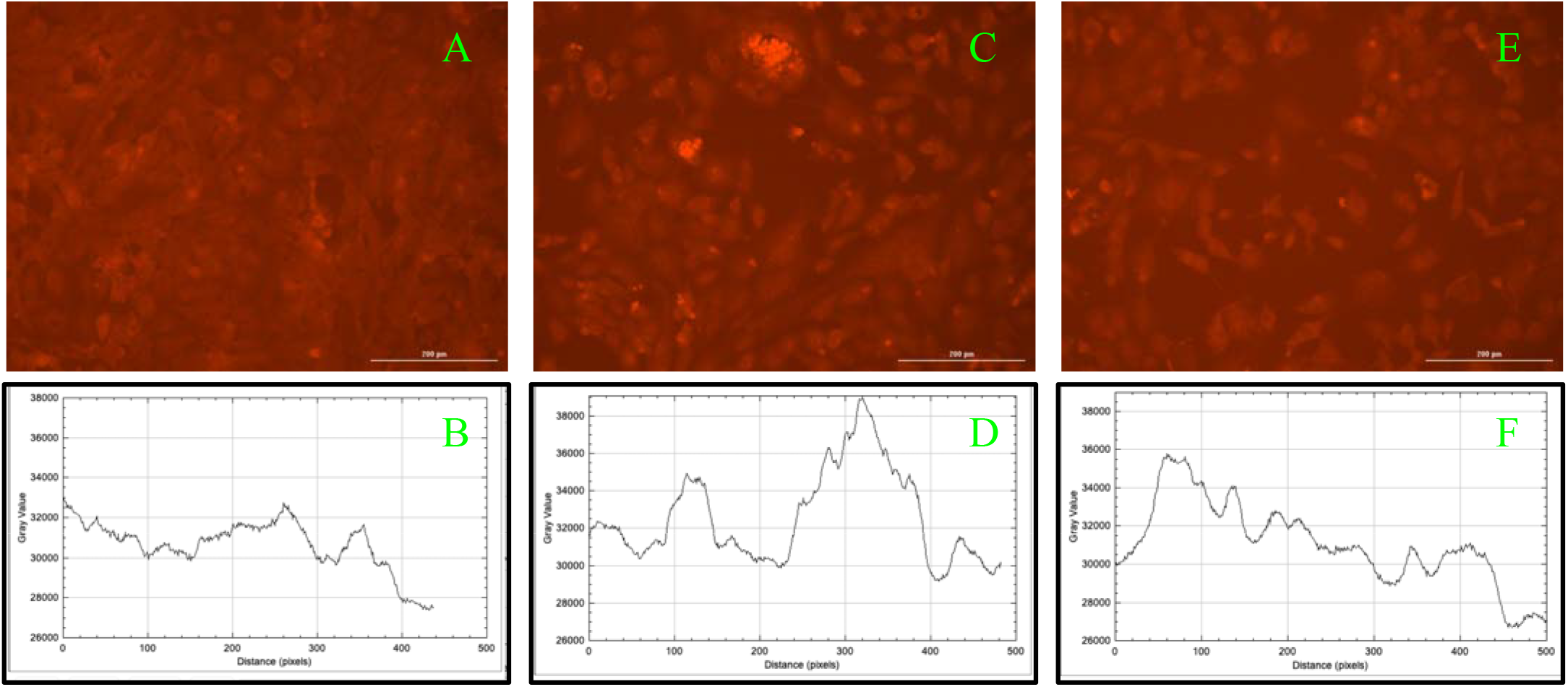
Light Intensity of β-Galactosidase between control and transfected astrocytes. Cells were imaged at 587nm under Texas Red (A,B,C) Images were converted to 8-bit grayscale and plot was produced representing mean gray value which corresponds to intensity of β-Galactosidase signal (B,D,F) (A) A representative image of Untreated Control cells is shown. (B) Light intensity plot for image A. (C) Representative image of scrambled plasmid cells treated with D-galactose for 72hrs (D) Light intensity plot for image C with peaks corresponding to signals with higher β-Galactosidase staining (E) Representative image for shSCN9A cells treated with D-galactose for 72hrs (F) Light intensity plot for image C with peaks corresponding to β- Galactosidase staining. *Images taken in Biotek Gen 5 with plots developed in FIJI-ImageJ*

This novel role of SCN9A was further verified by quantifying key markers of senescence including SA-□*-*gal and γ-H2AX. At 48 and 72 hours, SA-□*-*gal was measured in astrocytes. Results support the notion that SCN9A was able to prevent the cell from expressing the signal as compared to the control and scrambled when exposed to D-galactose (**Figure 7**). As compared to the untreated control wells, the control wells treated with D-galactose exhibited higher expression of β*-Galactosidase* (p<.05) and as expected, the scrambled RNA plasmid cells treated with D-galactose also exhibited higher expression of β*-Galactosidase* (p < 0.01) (Figure 7).

**Figure 7.**
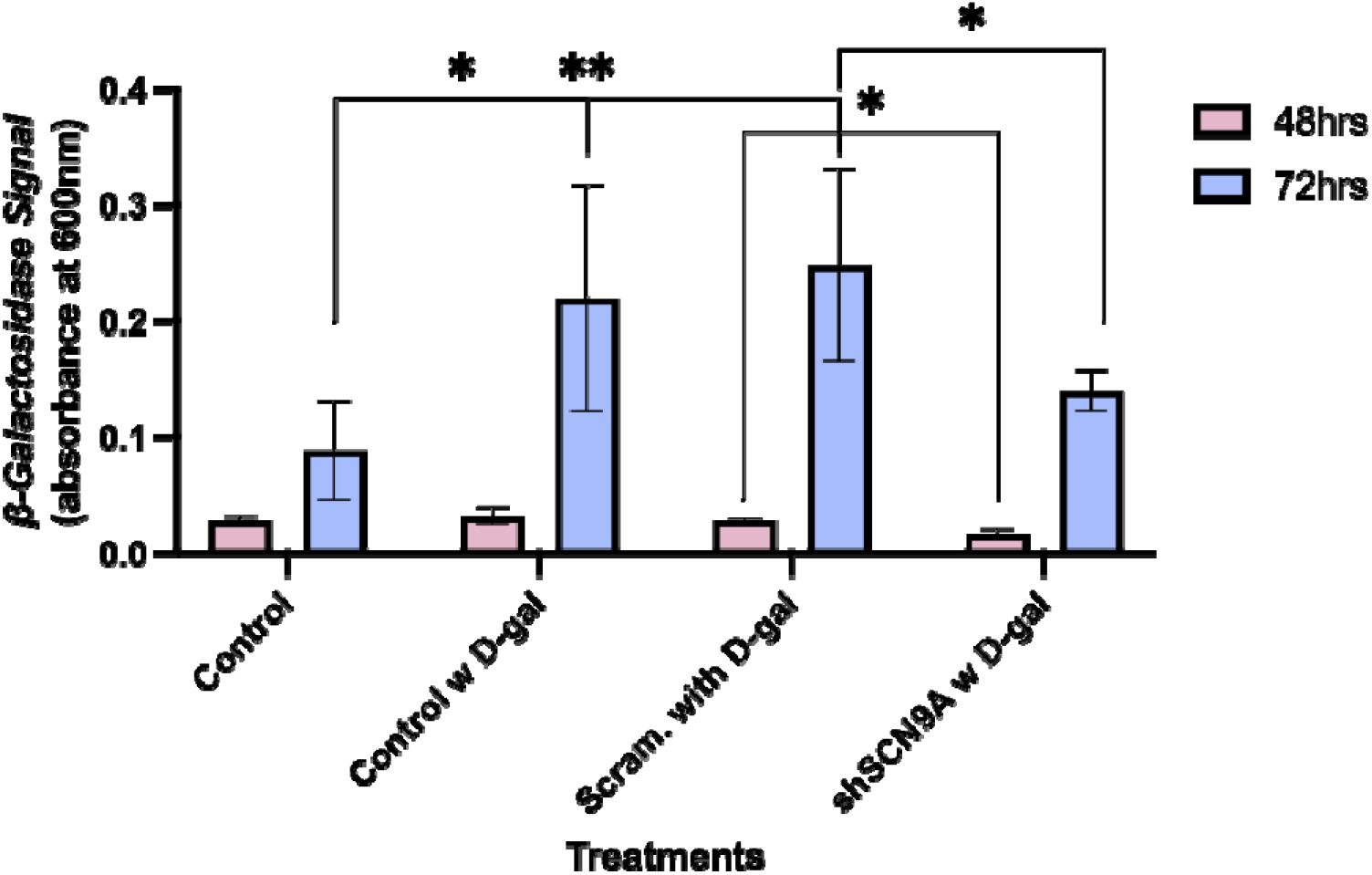
SCN9A knockout in Astrocytes prevented the development of β-Galactosidase after 48 and 72 hours of D-galactose (D-gal) exposure. Both Control and Scrambled RNA Astrocytes expressed higher levels of signal when measured at 600 nm as compared to the control group while the SCN9A knockout astrocytes did not reach these levels when exposed to D-galactose at both 72 and 48 hours (n=3). *p<.05 and **p<.01 ***p<.001 ****p<.0001

When comparing astrocytes with the shSCN9A plasmid and the scrambled plasmid, the cells with shSCN9A had a lower β*-Galactosidase* signal after 72 hours of D-galactose treatment which supports the hypothesis that inhibition of SCN9A will help prevent astrocytes from entering senescence (Figure 7). However, comparison of astrocytes with shSCN9A to control astrocytes showed a non-significant increase in signal, suggesting that the knockdown of SCN9A may not prevent all cells from entering senescence which is an interesting finding that should be investigated further.

Another marker of senescence is γ-H2AX; the phosphorylated form of Histone H2AX. The marker is sensitive to DNA damage and other common cellular signals of aging. It is notably considered the second most sensitive marker of senescence.^[16]^ Cells were exposed to D-galactose for 72 hours and relative γ-H2AX concentration was measured (**Figure 8**). The increased levels of γ-H2AX in cells exposed to D-galactose is consistent with the understanding that these cells have become senescent. However, if the knockout of SCN9A inhibited excess phosphorylation of Histone H2AX to γ-H2AX, the role of SCN9A in preventing senescence may be more intricate than initially anticipated.

**Figure 8.**
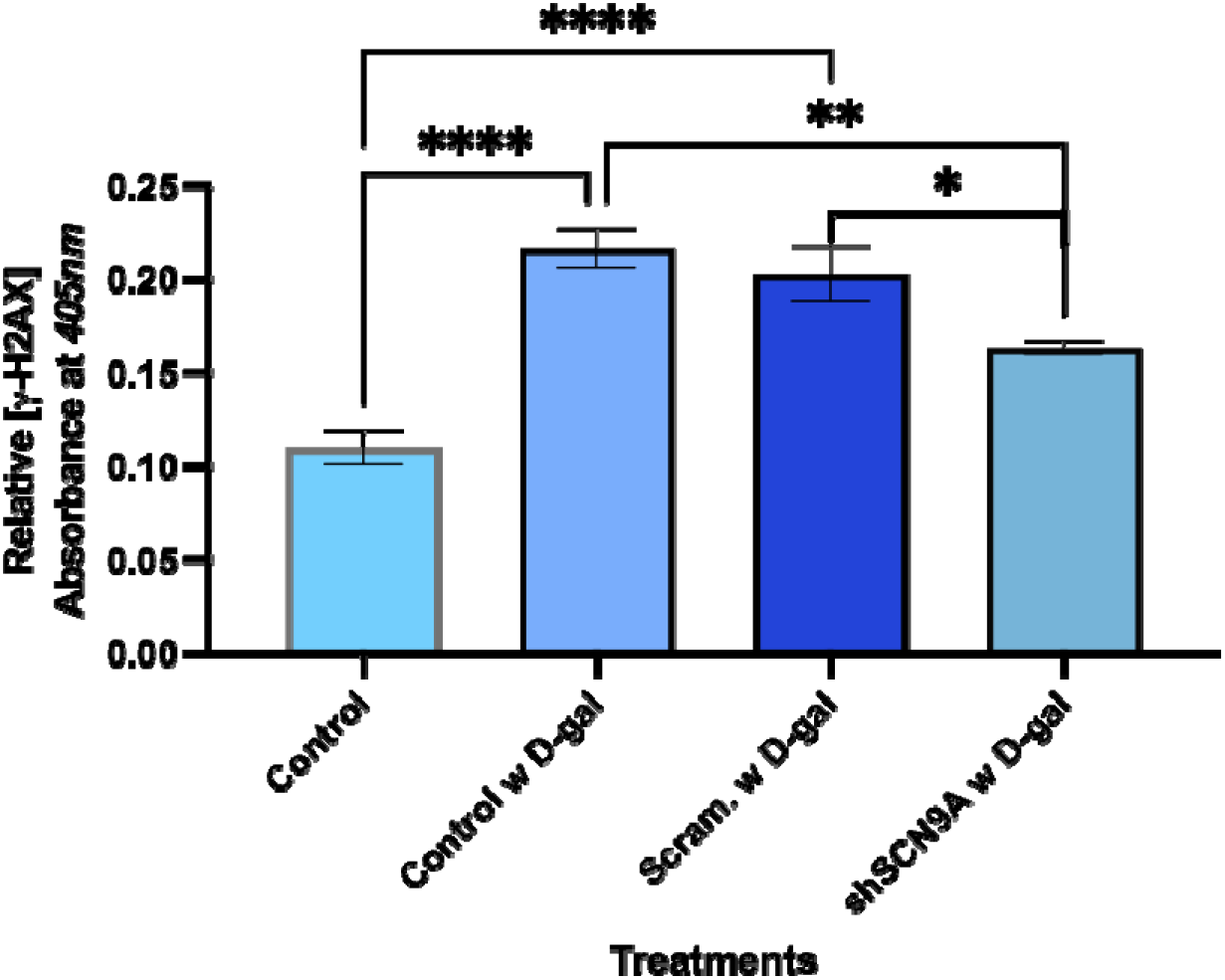
SCN9A knockout prevents astrocytes from excess phosphorylation of H2AX after 72 hrs of D-galactose (D-gal) exposure (n=3). *p<.05 and **p<.01 ***p<.001 ****p<.0001

Damage-associated molecular pattern molecules (DAMPs) have been well associated with aging. The increased levels of DAMPs in aging cells could serve as a marker for senescence as well. One release method of DAMPs without cell lysis is by measuring cell-free DNA (cfDNA).^[17]^ Transfected and control cells were plated with cell count normalized and exposed to D-galactose. After 72 hours, cfDNA levels were measured (**Figure 9**).

**Figure 9.**
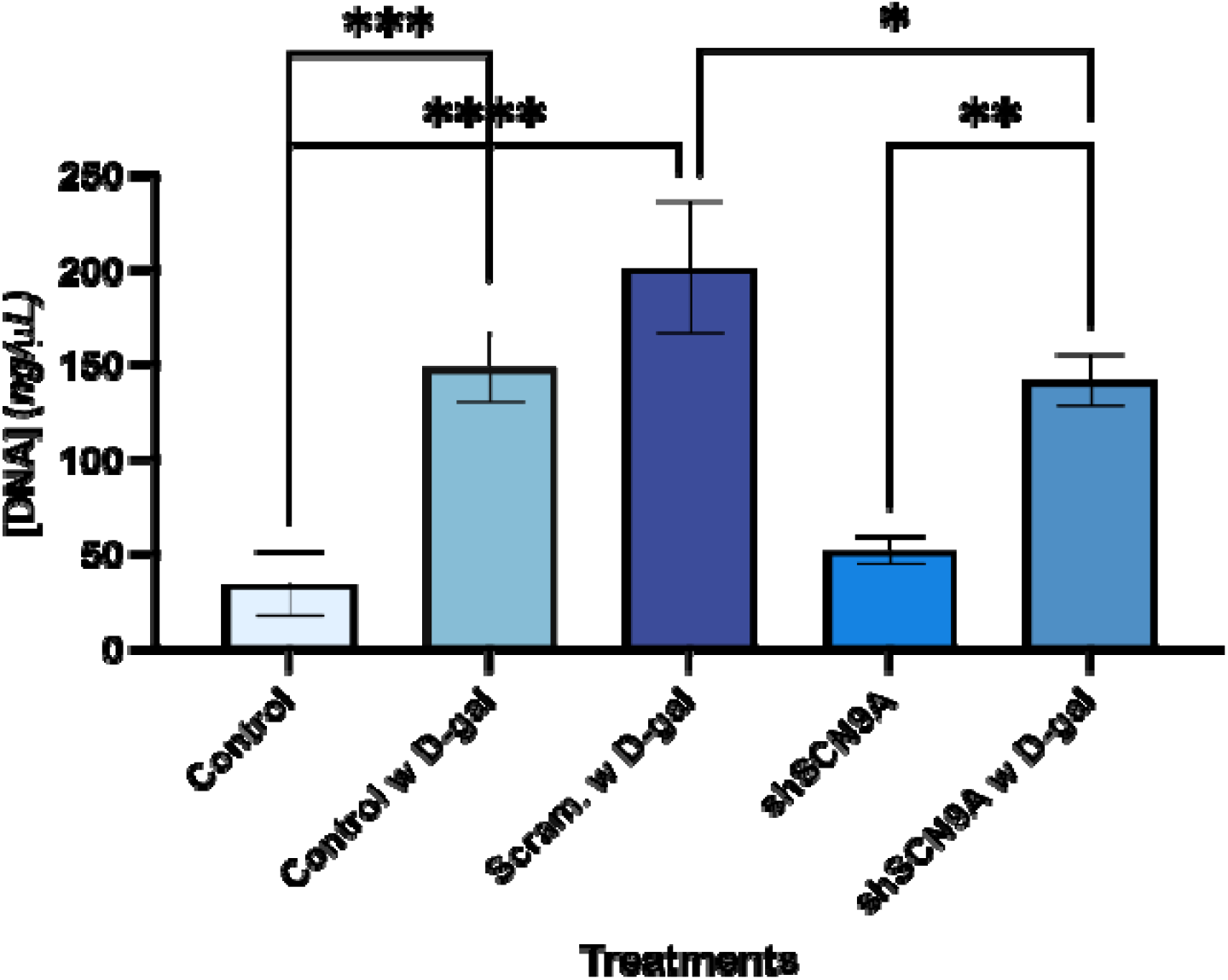
cfDNA levels are elevated in cells exposed to D-galactose after 72 hours (n=3). However, SCN9A inhibition prevented these elevated levels from rising as compared to cells with the Scrambled RNA. *p<.05 and **p<.01 ***p<.001 ****p<.0001

As expected, all cells exposed to D-galactose have higher levels of cfDNA suggesting DNA leakage as one of the consequences of the aggregation of DAMP. However, the cells with knockout of SCN9A exhibited lower levels of cfDNA suggesting that the mechanism in which SCN9A knockout inhibits senescence may also lead to the inhibition of the DAMP pathways. This result suggests a new avenue to explore regarding ion channel involvement in cell cycle pathways.

## 3. Conclusion

The results of this study reveal a novel indication that the knockdown of SCN9A, a gene that encodes for voltage-gated ion channel NaV1.7, has the ability to reverse senescence in astrocytes. While the gene has been shown as a pro-senescence gene in other cellular models, its role in the nervous system has yet to be discussed.

Investigations began with analysis of the key pathways responsible for aging in cortical neurons. The result of which suggested that ion channel and depolarization pathways were some of the most significant. In the nervous system, aging is associated with neuroinflammation including the presence of senescence-associated secretory phenotype (SASP) which includes the upregulation of IL-1 and IL-6. As neuroinflammation and ion channels were significant in the first dataset, the results suggest a link between ion channel dysfunction and excess neuroinflammation in patient’s brains with age. This also suggests a link between ion channel dysfunction and DAMP release as DAMP release’s correlation with neuroinflammation is well known.

To confirm that SCN9A could also play a role in the nervous system, pathway analysis was performed again which revealed that SCN9A knockdown in epithelial cells were associated with the same pathways that led to senescence in astrocytes. This showed that the senescence pathway in astrocytes follows a similar pattern as that in epithelial cells as both require a stress factor to induce a change in cell cycle regulation which will thus lead the cell to enter proliferation arrest.

Pathway analysis revealed that SCN9A expression in combination with an external stress factor can lead to ion channel dysfunction. Consequently, the senescence pathway is activated in cells which causes issues corresponding to DNA replication and mitosis. Thus, by inhibiting SCN9A expression when astrocytes are experiencing an external stress factor, it may be possible to prevent the cells from entering proliferation arrest *in vitro*. When exposed to D-galactose, a known inducer of senescence, transfected shSCN9A cells seemed to be able to avoid entering proliferation arrest and thus continued to enter the cell replication cycle normally as noted by confluency after treatment. In addition when staining cells for β*-Galactosidase,* the most accepted marker of senescence, cells containing the shSCN9A plasmid were more resistant to the stress of the D-galactose with less β*-Galactosidase* activity noted after 48 and 72 hours. The same held true when measuring levels of γ-H2AX; shSCN9A astrocytes were able to prevent excess phosphorylation suggesting a novel role for the gene in preventing proliferation arrest. Most notably, this suggests the ability of SCN9A to serve as a “gatekeeper gene” in astrocyte senescence which has never been presented before.

As SCN9A encodes for an ion channel, the results from this study predict that its crucial role in preventing senescence must be through some involvement in the neuroinflammation pathway. This is proposed because *in silico* analysis showed a connection between ion channels and neuroinflammation in aging neurons which has also been previously suggested. To this date, SCN9A has never been flagged as a key gene in the senescence pathway and these results point to its role being more significant than previously suggested in literature. While *in vitro* results showed that shSCN9A cells did not completely prevent all cells from entering senescence, its ability to significantly decrease the number of senescent cells as compared to the control is an undiscovered conclusion. Thus, inhibition of SCN9A expression could allow for an improvement in brain function by preventing astrocyte senescence. Going further, astrocyte senescence has implications with neurodegenerative diseases, specifically Alzheimer’s Disease. If SCN9A had potential to inhibit senescence *in vitro*, it may have some role in preventing the development of Alzheimer’s. Further research will be essential to understand the full role of SCN9A in neurodegeneration and how it may be used as a tool to treat neurodegenerative disorders.

## 4. Experimental Section/Methods

### In Silico Gene Expression Analysis of RNAseq data

Gene expression data (GSE53890) of cortical neurons between patients (n = 41) of age 21-103 were split into two cohorts: 20 patients aged less than 65 and 21 patients aged greater than 65. Cutoff age was established at 65 as this is the age when the incidence of AD diagnosis increases. GEO2R, an interactive web tool that uses GEOquery and limma R packages from the Bioconductor project, identifies differentially expressed genes (DEGs) between the two cohorts. STRING, a database that identifies known and hypothesized protein-protein interactions, was used to identify interactions and pathways between the differentially expressed genes.

Gene expression profile data (GSE110884) of immortalized mammary epithelial cells (n = 6) that underwent treatment of 4-hydroxytamoxifen to induce senescence was split into two groups: a treated group of 3 wells of epithelial cells transfected with an SCN9A shRNA and a control group of 3 wells of epithelial cells. GEO2R was used to identify differentially expressed genes (DEGs) between the two groups. Gene Ontology (GO) Analysis conducted through Enrichr identified significant pathways associated with the differentially expressed genes.

#### Cell Culture

Mouse (*Mus musculus)* astrocyte type III clones (C8-D30) were obtained from American Type Culture Collection (ATCC, Manassas, VA). Cells were cultured at 37L and 5% CO_2_ and maintained in Dulbecco’s Modified Eagle Medium (DMEM) (Invitrogen, Grand Island, NY) supplemented with 10% heat-inactivated fetal bovine serum (Invitrogen), 1% Penicillin/Streptomycin (Invitrogen), and 1% N2 growth supplement (Invitrogen). Upon confluency, cells were subcultured by trypsinization with 0.05% trypsin-EDTA (Invitrogen). Cell density was measured by trypan blue exclusion with a Vi-CELL XR Cell Viability Analyzer (Beckman Coulter, Indianapolis, IN).

#### D-galactose treatment and Cell Viability Assay

C8-D30 were treated with D-Galactose (Sigma-Aldrich CAS: 59-23-4) dissolved in DMEM at concentrations of 100 mM, 200 mM, 250 mM, 300 mM, and 555 mM for 48 hrs; untreated cells were used as control. An MTS-based CellTiter® 96 AQueous One (Promega, Madison, WI) assay was then used to measure cell proliferation and determine median inhibitory concentration (IC50) using a linear regression. After 48 hrs, treatment media was aspirated and 15 μL of Cell Titer reagent was added to each well followed by incubation for 1 h at 37°C and 5% CO_2_. After incubation, absorbance was measured at 490 nm with a BioTek ELX808 microplate spectrophotometer; A_490nm_ is directly proportional to the number of living cells in the sample. Untreated control cells were set as 100% viability and absorbance values converted to percent of control.

#### SCN9A Knockdown by RNAi

A kill curve, in which C8-D30 cells were treated with varying concentrations the selection antibiotic puromycin (0-10 µg/mL, Invitrogen) and monitored by light microscopy over 7 days for cell death, was used to determine the optimal selection concentration. The SCN9A gene was knocked down by transfecting cells with a pGFP-V-RS vector containing a short-hairpin RNA (shRNA) coding sequence complementary to SCN9A mRNA, or a scrambled control sequence (OriGene). C8-D30 cells were plated in a 24- well plate at a density of .5×10^5^ cells per well, in triplicate for each construct, in complete growth medium the day prior to transfection. Immediately before transfection, 1 µg plasmid DNA was diluted in 250 µL serum free medium with gentle vortex mixing. DNA complexes were prepared by adding 4 µL of TurboFectin to the diluted DNA and mixed by gentle pipetting; this solution was incubated for 15 minutes at room temperature, then added dropwise to the cells. GFP expression, visualized by fluorescence microscopy, was used as an indicator of plasmid uptake and only after 90% of cells visually showed successful knockdown of SCN9A were further experiments conducted. To create a stable SCN9A knock down lineage, cells were grown in complete media plus 6 µg/mL puromycin plate to maintain selective pressure, After passing cells into a T-25 flask, the transfected cells were grown out and used for further experimentation.

#### cfDNA measurement

Cell supernatant was collected from cell culture plates post 72- hour treatment. Each well was collected and stored individually. Cell culture media was used as a blank control to standardize any impurities found in the media. Using a Nanodrop Spectrophotometer (Thermofisher), DNA concentration was measured at 260 nm for each sample collected (ng/µL).

#### □-Galactosidase Assay

A □-Galactosidase-based histochemical staining kit was used to analyze senescence in C8D30 cells. Cells were plated at a concentration of 25,000 cells per well into a 96-well plate. After 24 hours for attachment, cells were treated with D-Galactose at IC50 of 250mM dissolved in complete or selection media; untreated cells were used as controls. After 72 or 48 hour incubation, culture media was aspirated and the cells washed with PBS (phosphate buffered saline). Cells were fixed for 10 minutes followed by 3 PBS washes. Staining solution was added to each well and the plate incubated overnight at 37°C without CO_2_. After incubation, the staining solution was replaced with PBS for imaging with a Cytation 3 multimode imaging system and *Gen*5 Data Analysis software (BioTek). A random selection of images were recorded from each well using a 10X and 60X objective, in brightfield and with a Texas Red filter (586 nm). Representative images were selected for further analysis in ImageJ to develop light intensity plots for gray-scale images in order to quantify β*-Galactosidase* staining.

#### γ-H2AX Assay

An enzyme linked immunosorbent assay, ELISA, was used to determine the relative expression of γ*-H2AX*. C8D30 cells post-72hr treatment were lysed using (Ethylenediaminetetraacetic acid)EDTA. Lysates were diluted in a coating buffer and added to a 96-well plate, in triplicate. After overnight incubation at 4°C, the plate was blocked with a buffer containing 1% bovine serum albumin (BSA) and incubated for 1 hour. The blocking solution was then replaced with the primary antibody, rabbit anti-mouse H2AX Monoclonal (VWR), diluted in the blocking buffer and the plate incubated once more. The secondary antibody, horseradish peroxidase-conjugated anti-rabbit IgG, diluted in blocking buffer, was added to each well and incubated. Secondary antibody solution was then removed and the plate washed with a buffer. The Horseradish peroxidase (HRP) substrate was added and absorbance measured 450 nm with a microplate spectrophotometer.

#### Statistical Analysis

Assays were repeated three or more times with 3 replicates (n = 3). Mean +/- standard deviations are reported in bar graphs unless otherwise noted. A Student’s t-test (unpaired, two-tailed) was conducted to determine statistical significance with an α-value of 0.05 unless otherwise noted. *p<.05 and **p<.01 ***p<.001 ****p<.0001

## Acknowledgments

We thank the Bergen County Academies for providing the resources to carry out the *in vitro* experiments in this study.

